# The interplay between facilitation and habitat type drive spatial vegetation patterns in global drylands

**DOI:** 10.1101/227330

**Authors:** Miguel Berdugo, Santiago Soliveres, Sonia Kéfi, Fernando T. Maestre

**Affiliations:** Departamento de Biología y Geología, Física y Química Inorgánica, Escuela Superior de Ciencias Experimentales y Tecnología, Universidad Rey Juan Carlos, C/ Tulipán s/n, 28933 Móstoles, Spain.; Institute of Plant Sciences, University of Bern, Altenbergrain 21, 3013 Bern, Switzerland.; ISEM, CNRS, Université de Montpellier, IRD, EPHE, Montpellier, France.

**Keywords:** Aridity, functional traits, grasslands, patch-size distributions, power laws, shrublands, spatial patterns

## Abstract

The size distribution of discrete plant patches (PSD), a common descriptor of the spatial patterns of vascular vegetation, has been linked to variations in land degradation and ecosystem functioning in drylands. However, most studies on PSDs conducted to date have focused on a single or a few study sites within a particular region. Therefore, little is know on the general typology and distribution of PSDs at the global scale, and on the relative importance of biotic and abiotic factors as drivers of their variation across geographical regions and habitat types. We analyzed 115 dryland plant communities from all continents except Antarctica to investigate the general typology of PSDs, and to assess the relative importance of biotic (plant cover, frequency of facilitation, soil amelioration, height of the dominant species) and abiotic (aridity and sand content) factors as drivers of PSDs across contrasting habitat types (shrublands and grasslands). We found that both power-law and lognormal PSDs were generally distributed regardless of the region of the world considered. The percentage of facilitated species in the community drives the emergence of power-law like spatial patterns in both shrublands and grasslands, although mediated by different mechanisms (soil and climatic amelioration, respectively). Other drivers of PSDs were habitat-specific: height of the dominant species and total cover were particularly strong drivers in shrublands and grasslands, respectively. The importance of biotic attributes as drivers of PSDs declined under the most arid conditions in both habitats. We observed that PSDs deviated from power law functions not only due to the loss of large, but also of small patches. Our results expand our knowledge about patch formation in drylands and the habitat-dependency of their drivers. They also highlight different ways in which facilitation may act on ecosystem functioning through the formation of plant spatial patterns.

## INTRODUCTION

Vegetation in arid, semi-arid and dry-subhumid ecosystems (drylands, hereafter) is usually arranged in a two-phase mosaic formed by plant patches interspersed in a matrix of open areas devoid of perennial vascular vegetation (Tongway et al. 2001). The frequency of size classes of these plant patches (patch-size distributions, referred to as PSD hereafter) can be characterized by heavy-tail distributions, which have been linked to changes in ecosystem functioning (Maestre and Escudero 2009, Berdugo et al. 2017b) and degradation status (Kéfi et al. 2007a, Scanlon et al. 2007) in drylands. Mathematical models suggest that PSDs of dryland vegetation follow power law functions (i.e., there are many small patches and a few very large ones, (Kéfi et al. 2007a, Scanlon et al. 2007), and that increasing environmental harshness (e.g. higher aridity or grazing pressure) reduces the number of large patches, thereby generating truncated power laws (Kéfi et al. 2007a, Lin et al. 2010).

The shift from power-law to other PSDs has been observed in field sites undergoing degradation (Kéfi et al. 2007a, Scanlon et al. 2007, Lin et al. 2010), but truncated power laws or lognormal functions have also been found to best fit PSDs in well preserved ecosystems (Maestre and Escudero 2009, von Hardenberg et al. 2010, Berdugo et al. 2017b). To explain these contrasting results, we need to better understand the ecological mechanisms by which PSDs shift from power law to other distributions. Plausible ecological mechanisms, such as the loss of large patches or the emergence of dominant scales (i.e., overrepresentation of some patch sizes) have been proposed to explain changes in PSDs (e.g., see Von Hardenberg et al. 2001, Xu et al. 2015). Nevertheless, we still lack empirical evaluations of the importance of these ecological processes in real-world ecosystems, which precludes a complete interpretation of the effects that plant spatial patterns may have on ecosystem functioning.

Facilitative interactions are often invoked as a major driver of PSDs in drylands, as they promote the formation of large patches that underpin the creation of power-law PSDs (Kéfi et al. 2007a, Scanlon et al. 2007, Xu et al. 2015, Berdugo et al. 2017b). Facilitative mechanisms that can increase the size of plant patches include improvements in soil and microclimatic conditions beneath nurses, which allow protegeé plants to thrive under environments to which they are poorly adapted (Maestre et al. 2003, Liancourt et al. 2017, Berdugo et al. 2017c). Whereas soil amelioration depends on the attributes of nurses (Pugnaire et al. 1996, Maestre et al. 2001) and the environmental conditions in which they live (e.g., sandy soils exhibit the largest differences in fertility between nurse and open areas, see Ochoa-Hueso et al. 2017), the effect of shading may be more influenced by the pool of beneficiary species and their relative adaptation to climate (Soliveres and Maestre 2014, Liancourt et al. 2017, Berdugo et al. 2017c). Both facilitative mechanisms could be important mediators of the impact of facilitation on ecosystem functioning by creating particular spatial patterns, a link that is still poorly understood (Cardinale et al. 2002, Maestre et al. 2010, Wright et al. 2017). Soil amelioration directly impacts average ecosystem functioning, and also increases soil heterogeneity (Dean et al. 1999, Ochoa-Hueso et al. 2017). On the other hand, preserving a high diversity of species may promote ecosystem functioning through niche complementarity (Tilman et al. 1997, Gross et al. 2017). Thus, disentangling the relative importance of different facilitative mechanisms on spatial pattern formation may inform us about how facilitation affects ecosystem functioning via complementary processes (spatial pattern formation versus direct soil amelioration and increased species richness).

PSDs are not only affected by degradation processes or facilitation. Functional traits of the dominant species (Aguiar and Sala 1999, Maestre and Cortina 2005, Borthagaray et al. 2012), soil attributes (Von Hardenberg et al. 2001) or plant cover (Maestre and Escudero 2009) can also importantly affect PSDs. For instance, high cover might promote vegetation clumping due to lack of space (Abades et al. 2014, Xu et al. 2015), while sandy soils usually alter the distribution of resources causing interactions with facilitation mechanisms such as differential infiltration rates (Rietkerk et al. 2004). Also, tall species may influence the basal patch sizes from which PSDs emerge. Importantly, the factors driving PSDs may differ across habitat types (Goslee et al. 2003, Lett and Knapp 2003, Bordeu et al. 2016), although this is poorly understood due to the lack of cross-habitat studies. Contrasting dominant plant types, such as grasses versus shrubs, exhibit different ways of reproduction (e.g. grasses tend to reproduce via rhizomatous roots) and resource uptake (grasses have shallower roots than shrubs) that inherently affect spatial pattern formation (Goslee et al. 2003, Lett and Knapp 2003, Ravi et al. 2008). Also, grasses and shrubs differentially affect ecohydrological mechanisms such as run-off, erosion or evapotranspiration, with shrubs producing higher differences in water availability with adjacent bare ground areas than grasses (Huxman et al. 2005, Ludwig et al. 2005, Okin et al. 2009, Berdugo et al. 2014). These ecohydrological effects have been previously documented to affect the way in which grasses and shrubs interact with other species (Aguiar et al. 1992, Maestre et al. 2003). Therefore, differences between habitat types could also influence how processes such as plant-plant interactions affect PSDs. Increases in aridity have been found to be correlated with reduced vegetation cover (Delgado-Baquerizo et al. 2013) and different relative dominance of grasses vs. shrubs in drylands (Knapp et al. 2008). Therefore, understanding the interactions between aridity and other PSD drivers is of paramount importance to better understand and forecast how dryland ecosystems respond to ongoing climate change.

The current lack of understanding of the relative importance of biotic vs abiotic drivers of PSDs in drylands restricts our ability to forecast changes in their structure in response to climate change, and to use them as indicators of land degradation in these areas. To contribute to filling this gap in our knowledge, we investigated the PSDs of perennial vegetation in 115 drylands from all continents except Antarctica spanning a wide range of environmental conditions, soil and vegetation types. This allowed us to assess: (i) the region and habitat-type dependency of PSDs in drylands worldwide, (ii) the relative importance of aridity and biotic (plant cover, facilitative interactions, soil amelioration, habitat type and plant functional traits) factors as drivers of PSDs, and (iii) whether the importance of different PSDs drivers changes depending on the habitat considered (shrublands vs. grasslands).

## MATERIAL AND METHODS

### Study sites and data collection

We studied 115 dryland ecosystems from 13 countries (Fig. 1), which are a subset of the 236 sites from Ochoa-Hueso et al. (2017). Annual mean temperature and rainfall ranged from 2.6 to 25.7 °C and from 67 to 801 mm, respectively. Elevation, latitude and longitude varied from 76 to 4524 m.a.s.l., from -41° to 40°, and from -115° to 142°, respectively. The database includes three habitat types (grasslands, shrublands and open woodlands/savanna); however, we did not use open woodlands/savanna because there were very few sites suitable for our analyses (see details below). Grassland and shrubland sites covered a wide range in species richness (2 to 39 perennial species) and total plant cover (from 4.5 to 82.8 %). See Maestre et al. (2012), for full details.

**Figure 1.**
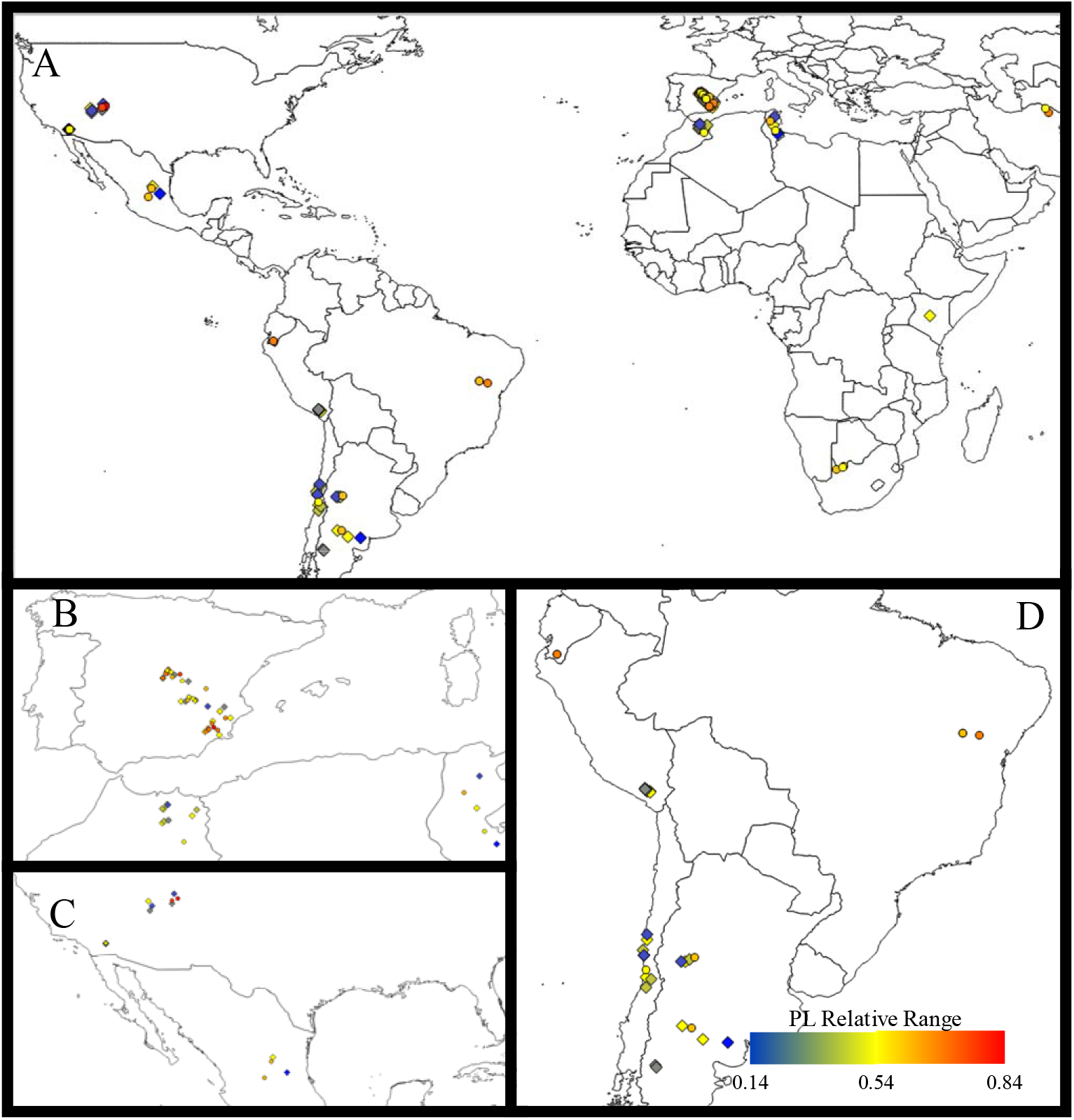
Map of patch-size distributions across the studied sites. The colour switch from blue to red according to the power law relative range value. A: Map of the world; B: zoom on Mediterranean area; C: Zoom on North America area; D: Zoom on South America area.

At each site we performed a standardized field survey protocol. We established four 30 m long transects separated 8 m from each other and measured the cover of perennial plants using the line intercept method as detailed in Maestre et al. (2012). We also collected five soil samples at 0-5 cm depth under the dominant species and in bare ground areas close by. These soils were sieved (2 mm mesh), air dried for one month and stored for laboratory analyses. To standardize soil analyses, the samples were shipped to Spain, where they were analyzed in the same laboratory.

### Measurement of patch-size distributions

From the 236 sites available, we used for this study 115 sites from which we were able to quantify their PSDs using high-resolution remote sensing data (see Berdugo et al., 2017b) for details). In brief, we downloaded VirtualEarth (http://www.bing.com/maps) and Google Earth (https://earth.google.com) images with resolution ≤ 30cm/pixel. To obtain a sufficient number of patches as to fit power law functions to their PSDs, we took three adjacent subplots of 50m x 50m per site. The first one was placed to fit that surveyed in the field. We classified the images by using authomatic luminance threshold detection, and contrasted their results with those from other classification methods based on expert knowledge (see Appendix 1 for details). Finally, remote-sensing cover estimates from the first subplot were correlated with field measurements to ensure that the classification procedure reproduced what was observed on the field. The image classification analyses used fitted reasonably well observed field data (see Appendix, Fig. S1).

For each site, we extracted all the patches and their sizes resulting from the image classification analyses of the three subplots. We pooled them and fitted a power law to their distributions. We used the approach of Clauset et al. (2009) to get the two main parameters of power law distributions:

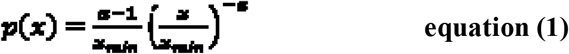

where x represents the patch size and p(x) describes the frequency of patches of a certain size. This equation represents the probability density function of a power decay. The parameters of the distribution are x_min_, the minimum patch size from which the fit to a power law starts (below that point data are discarded from the fitting procedure) and α, the rate of decay of frequency with patch sizes (see details in Berdugo et al., 2017b).

Sometimes the range of data remaining after discarding patch sizes lower than x_min_ is not representative of the observed PSD, especially when it is curved and best fits a lognormal function. To obtain an estimation of the range of patch sizes to which a power law could be fitted, we calculated the power law relative range (PLR) as:

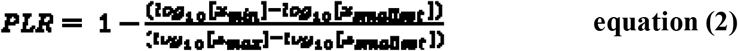

where, x_smallest_ and x_max_ are the size of the smallest and largest patch in the image, respectively. PLR theoretically varies from 1 (all data fitted to a power law function) to 0 (no data fits a power law function). PLR is related to the shape of the distribution and thereby to the goodness of fit to a power law (Berdugo et al. 2017b), but is not exclusive to power law distributions (i.e. may be used for other heavy tailed distributions as well; e.g. simulated lognormal distributions fitted using this methodology had a PLR around 0.3-0.4). The use of the PLR allowed us to: i) compare all PSDs among our sites, which vary from power law to lognormal, using a standard methodology for all of them, and ii) produce general descriptors of all the PSDs evaluated, independent of whether they fitted better to a power law or a lognormal function.

### Assessment of biotic and environmental factors

At each site, we measured biotic and environmental attributes known to influence PSDs: sand content, aridity, plant cover, height of the dominant species, percentage of facilitated species and soil amelioration (the increase in soil fertility under plant patches). Further rationale for the selection of these variables is presented in Appendix 2.

We measured sand content according to Kettler et al. (2001). The aridity level for each site was calculated as 1 – Aridity Index (AI). AI is precipitation/potential evapotranspiration, obtained from Zomer et al. (2008), which use climatic interpolations provided by Worldclim (Hijmans et al. 2005). Total perennial cover was estimated using the line-intercept method along four 30-m long transects within each site, and ranged from 4% to 83% in our study sites. The height of the dominant species was obtained from previous studies, local floras and global databases (Kattge et al., 2011; see Soliveres et al., 2014, Le Bagousse-Pinguet et al. 2017, for more details on trait data acquisition). We introduced habitat type (grassland and shrubland) to control for differences in the growth form of the dominant species. Habitat types were identified both types depending on the dominant plant form inhabiting the sites.

We measured plant-plant interactions in two ways to disentangle their potential effects on PSDs mediated either by their frequency or the strength of soil amelioration. First, as a measure of the frequency of positive plant-plant interactions, we quantified the proportion of species more associated with a given nurse than expected by chance. For doing so, we compared the number of individuals found in open sites vs. those found under nurses and calculated a χ^2^ value for each pairwise interaction (see Soliveres et al., 2014 for full details). The frequency of facilitative interactions was measured in a subset of 70 sites. Second, the strength of soil amelioration may increase the survival and growth of beneficiaries by nurses. We assessed it by measuring the difference in the organic carbon contents obtained in vegetated and bare ground areas (Allington and Valone 2014). We measured organic carbon by colorimetry after oxidation with a mixture of potassium dichromate and sulfuric acid (Anderson and Ingram 1994).

## Statistical analyses

### Describing the general typology of patch-size distributions

The way in which PLR decrease may indicate processes of interest (e.g., decreases in PLR associated with less small patches would indicate lack of recruitment, whereas a lack of large patches would impede the clustering of plant patches). Thus, we attempted to explore how the geometry of PSDs changed with decreasing PLR. The parameters x_max_ and x_min_ (corresponding to the maximum patch size and the patch size of the initialization point of a power law, respectively) are involved in the calculation of PLR. Hence, according to eqn 2, we expect PLR to decrease with higher x_min_ or lower x_max_. By relating both parameters to PLR we expect to gain insights into how PLR shortening occurs in curved PSDs. It might be either that: i) large patches become more infrequent with departures from power law fitting, thus shrinking PLR from the right side of the PSD; ii) the size of the large patches remain constant in all the study sites and that PLR is reduced because power law initializes in larger patch sizes (thus x_m_;_n_ increases while x_max_ remain constant; iii) that PLR is diminished as a consequence of both processes at the same time. To visualize the change in overall geometry of PSDs with decreasing PLR, we collapsed all PSDs into a single plot.

### Identifying the drivers of patch-size distributions

Prior to analyze the drivers of PLR, we fitted a statistical model to PLR as a function of both latitude and longitude to assess whether PSDs exhibited any biogeographic trend. We also compared the PLR of different zones of the world to evaluate whether these biogeographic patterns were found in different study areas. To defined the zones as: North America; South America, Mediterranean basin; Asia and Australia.

We used multi-model inference to assess how environmental and biotic drivers affect observed PSDs. This analytical procedure fits all possible combinations of predictors and ranks the models obtained according to their Akaike information criterion, corrected for small sample size/ number of predictors ratio (AICc). Models deviating less than two units of AICc from the best model, i.e. that with the lowest AICc, are not considered different from it (Burnham and Anderson 2004). Between the best models selected according to this criterion, a weighted average of standardized effects for each predictor is calculated (see Lukacs et al., 2009). We built a model using PLR as a response variable, and sand content, aridity, habitat, cover, height of the dominant species, facilitation and soil amelioration as predictors. We evaluated all possible interactions of biotic attributes with aridity to assess whether the deviations of PSDs from power laws that has been hypothesized when aridity increases (Kéfi et al. 2007a) acts through interactions with biotic factors. To know whether different habitats differ on the relative importance of other predictors as drives of PLR, we introduced all possible habitat × rest of predictors interactions. We repeated the analysis for grasslands and shrublands separately to better show the interaction between habitat type and the other factors evaluated. We performed this analysis with those sites for which we had all the information (N=71), but analyses with the sites that did not include facilitation data (N = 111) were consistent with those shown here (see Appendix, Fig. S2).

We performed multi-model inference analysis with the package MuMIn (Barton 2016) in R (R Development Core Team 2008). Prior to model fitting, we box-cox transformed the response variable to approximate normality of the residuals using the function boxcox of the package MASS (Venables and Ripley 2002) in R. Primary data are available in figshare <doi: 10.6084/m9.figshare.5640193> (Berdugo et al. 2017a).

## RESULTS

The PSDs of all 115 sites were heavy-tailed with varying levels of curvature (Fig. 2). We found that the increase of x_min_ and the decrease of x_max_ acted in tandem to create deviations of PSDs from a power law function (Appendix Fig. S3). As PLR decreased, PSDs tend to curve by exhibiting a plateau in the first part of the distributions, thus suggesting the emergence of a predominant scale (Fig. 2). A few distributions with a very low PLR showed ample curves with high x_max_ values (Fig. 2i), indicating the absence of processes forming power laws. Some of the distributions with the highest PLR had spanning clusters (i.e., abnormally large patches that are usually formed by vegetation merging in high covered sites, see Abades et al. 2014, see Fig. 2v). In sum, the curvature of distributions with low PLR was explained mainly by a loss of large patches, but also by a decrease in the relative proportion of small patches compared to the distributions showing higher PLR (Fig. 2).

**Figure 2.**
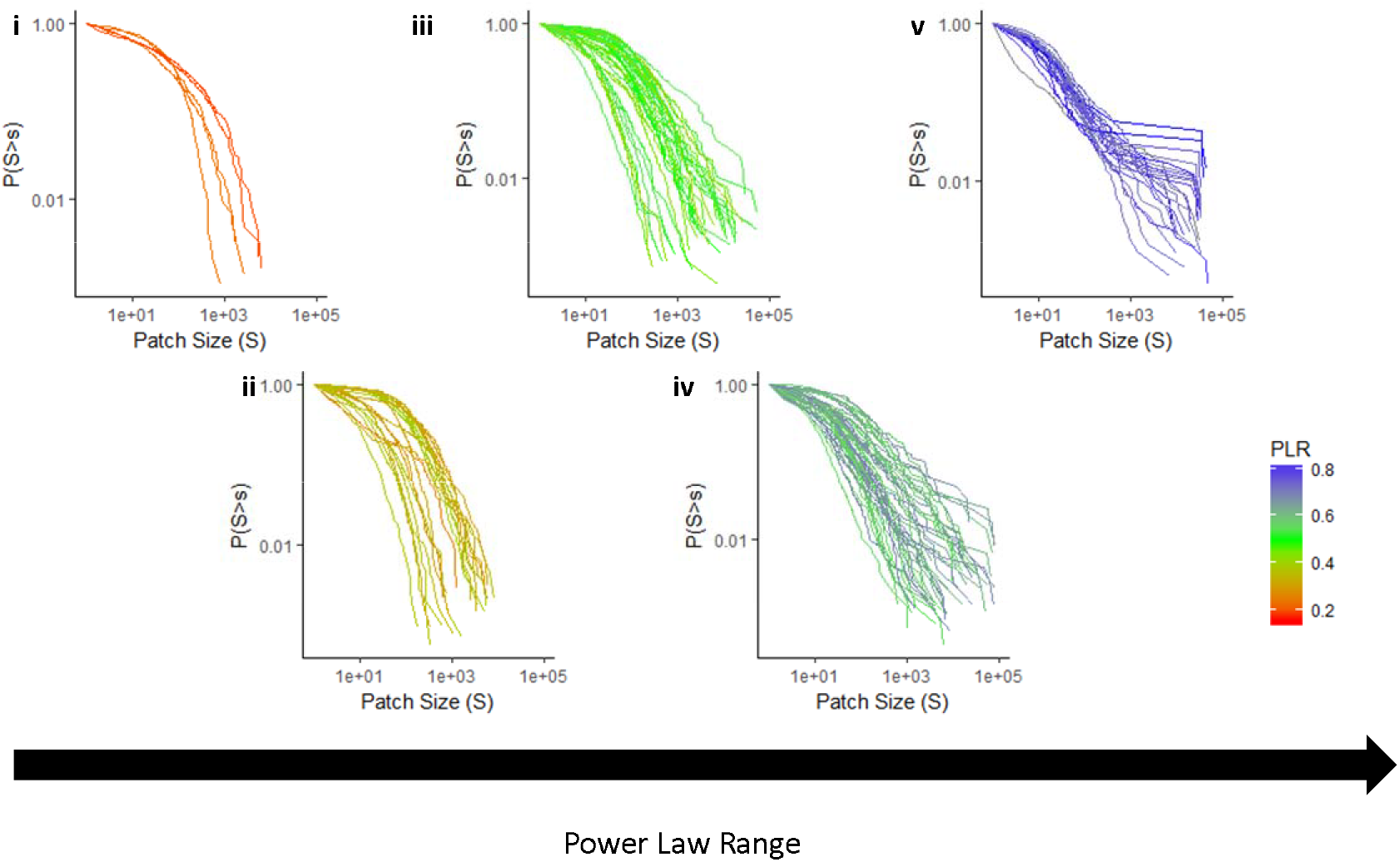
Patch-size distributions of perennial vegetation in the drylands studied. PSDs with increasing PLR from left to right (i: 0.1-0.25; ii: 0.25-0.4; iii: 0.4-0.55; iv: 0.55-0.7 and v: 0.7-0.85). Scales of both x and y axis are the same to ensure comparability. Color indicates the power law relative range of distributions (PLR, according to the legend), with higher values indicating a closer fit to a power law function.

PLR values did not vary with latitude, and increased slighty with longitude (Fig. 1). However, PLR maintained similar values across regions of the world (except for Africa and Asia, where they were higher compared to the Mediterranean basin and South America and North America, although a few points were sampled in the former regions, Appendix, Fig. S4). When we analyzed all habitat types together, total cover, percentage of facilitated species and the height of the dominant species were the main biotic factors driving the emergence of power laws in PSDs (Fig. 3). The two latter interacted significantly with aridity, with a reduced effect of plant height and facilitation under drier conditions. Aridity and sand content were only significant in the models with the highest statistical power (i.e., without percentage of facilitated species as a predictor, Appendix Fig. S2) and led in both cases to more curved PSDs.

**Figure 3.**
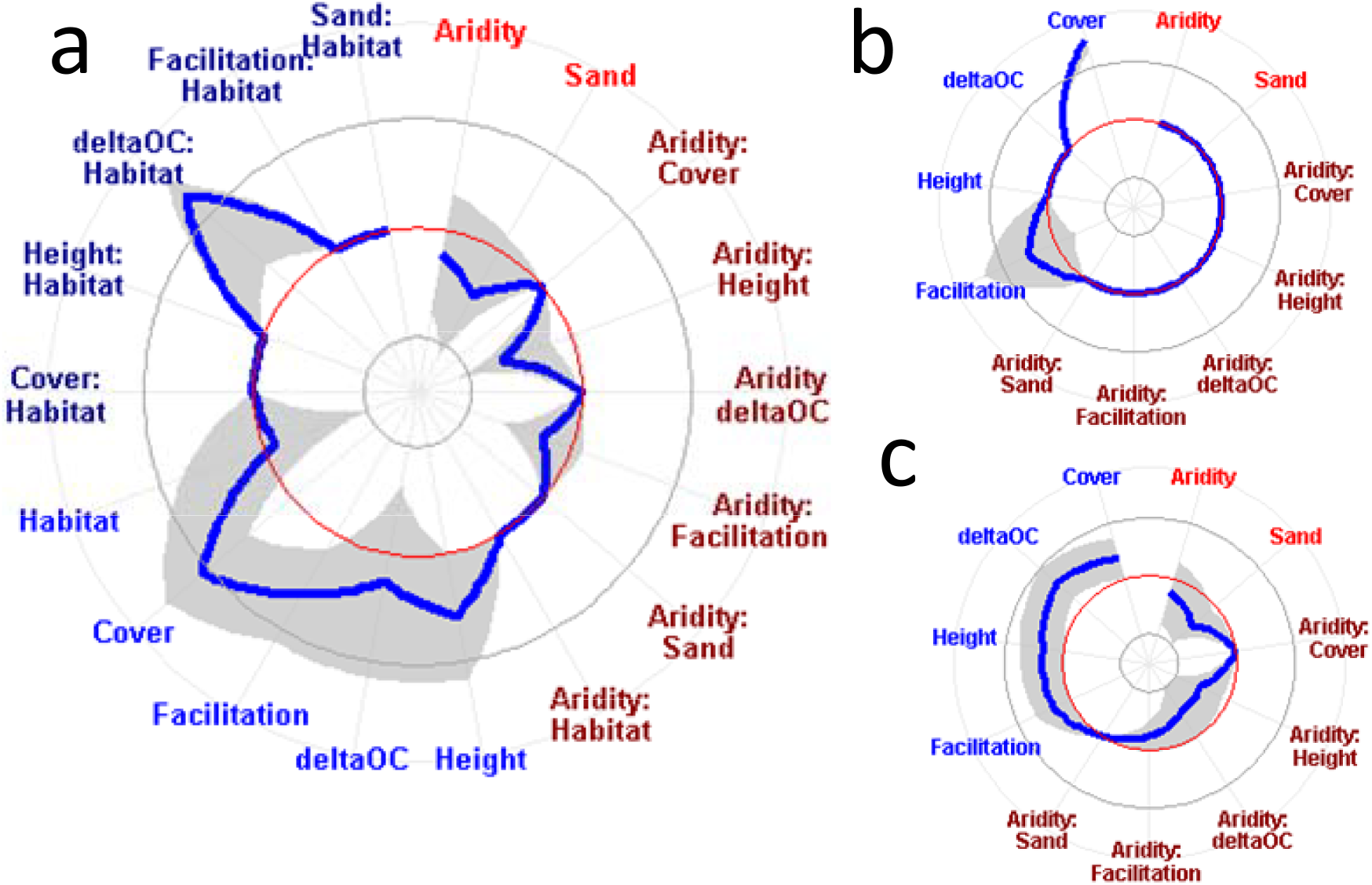
Drivers of patch-size distributions in global drylands. Standardized effect sizes of drivers of power law range for all habitat types (A; N= 70), grasslands (B; N = 37) and shrublands (C; N = 33) after model averaging of the best models in multimodel inference analysis (ΔAIC respect from the best model <2). Abiotic (red) and biotic (blue) drivers, and the interactions between them (dark red) and with habitat type (dark blue) are displayed. Shaded areas represent 95% confidence intervals of effect sizes; the red line represents effect sizes of 0 and the inner and outter dark grey lines represent effect sizes of -1 and 1, respectively.

The importance of PLR drivers varied with the habitat type considered, as suggested by the interactions between habitat type and soil amelioration in the overall model (Fig. 3a). In grasslands, the main predictor of PLR was total cover, followed by the percentage of facilitated species (Fig. 3b). In shrublands, height, percentage of facilitated species and soil amelioration all promoted the emergence of power laws in PSDs (Fig. 3c). Effects of total cover on the emergence of power laws were, however, dampened and even disappeared in the models with more statistical power in shrublands (Appendix, Fig. S2). Soil sand content was an important driver reducing the occurrence of large patches in shrublands, but not in grasslands. As with facilitation and plant height, increasing aridity also dampened the effects of soil amelioration on PLR.

## DISCUSSION

### Typology of patch-size distributions

Studies conducted over a few sites have shown that the PSDs of dryland vegetation can be fitted to a power law function (Scanlon et al. 2007, Lin et al. 2010, Moreno de las Heras et al. 2011). Our results confirmed these findings, and extended them to a wide variety of dryland communities worldwide. We also showed that variations in PSDs are not driven by geographical changes, but rather depend on the habitat type considered. The truncation of power law patch distributions has been associated to the loss of large patches in ecosystems undergoing degradation (Kéfi et al. 2007a, Lin et al. 2010). However, we also observed deviations from power law functions due to the lost of not only the large, but also the small patches. This is shown by the curvature observed both on the left and right parts of the PSD (Fig. 2). The loss of both large and small patches is not often invoked as a major driver of change in PSDs, although similar patterns to those observed here globally have been already found in the field (Quets et al. 2013, Berdugo et al. 2017b), as well as in theoretical studies (Marani et al. 2006, von Hardenberg et al. 2010). The loss of small patches could be explained by a reduction in the number of recruited seedlings under harsh environments (Weltzin and McPherson 1999), or by the lower number of isolated individuals found in ecosystems dominated by facilitation (Bertness and Callaway 1994, Soliveres et al. 2014).

Apart from this general pattern, we found several types of distributions in the studied drylands. For instance, some curved distributions (especially those with low PLR) appeared without showing a plateau before x_min_. These distributions probably are better fitted by exponential distributions than by lognormal ones. Both lognormal and exponential distributions have been fitted to PSDs (Manor and Shnerb 2008, von Hardenberg et al. 2010), but the former entails the emergence of a predominant scale, whereas exponential distributions do not. Importantly, PSDs fitted by exponential distributions may still contain large patches (Fig. 2i). This may be related with the overall scale of the system (i.e., when trees are present, patches can be large and not clumped, and thus their PSDs would not exhibit power laws). At the same time, we found evidence of the presence of spanning clusters (i.e., the appearance of very large patches spanning one side of the studied area to the other, see highest PLR cases in Fig. 2v). Spanning clusters are formed because high cover levels can increase the merging of vegetation when biomass reaches a saturating level, even if there is no mechanism promoting vegetation clumping (Abades et al. 2014, Xu et al. 2015). Although the formation of spanning clusters has been found to be more frequent at cover values around 60% in previous studies (Abades et al. 2014), we found these clusters in sites with lower cover values, while some sites with cover > 60% did not exhibit them (Appendix, Fig. S5).

### Abiotic factors drive the shape of patch-size distributions across habitats

Although aridity was not a significant driver of the change in PSDs, it strongly modulated the effects of biotic attributes on PSDs in shrublands, but not in grasslands. Grasslands in our dataset were particularly prevalent under moderate aridity conditions (Appendix, Fig. S6), so the lack of effect of aridity interactions might be a consequence of a lower aridity range in the grasslands surveyed. Our findings indicate that, in shrublands, aridity prevents biotic attributes (traits of the dominant species, plant-plant interactions) to form large patches under the most arid conditions. This provides empirical support to the often hypothesized facilitation collapse as the mechanism underlying shifts from power law to curved PSDs under extreme environments (Kéfi et al. 2007b). Some studies have observed or hypothesized a diminished importance of facilitation for community assembly as aridity increases (Holmgren and Scheffer 2010, Berdugo et al. 2017c). Although this pattern does not seem to hold for the effect of facilitation on species richness (Soliveres and Maestre 2014), unimodal facilitation-aridity relationships have been observed for the effect of facilitation on species abundances (Berdugo et al. 2017c). Such a decrease in the importance of facilitation has been related to community specialization to arid conditions (Berdugo et al. 2017c), and, therefore, may underpin the low importance of positive plant-plant interactions under these conditions. Our results link this decrease in facilitative interactions with the reduction in the frequency of large patches and to the abrupt changes in PSDs observed under extreme arid conditions (Berdugo et al. 2017b).

### Habitat-specific factors drive the shape of patch-size distributions

We found large differences in the drivers of PSDs depending on the habitat type considered. In grasslands, the main factor controlling the emergence of large patches was total cover. It is important to note that we cannot differentiate cause from consequence (cover driving spatial pattern or the other way around) from our observational study. On the one hand, some studies have linked patch formation with increases in the ability of plants to maintain a high biomass due to an increased resource capture efficiency (Aguiar and Sala 1999, Boer and Puigdefábregas 2005). On the other hand, as we already discussed, high cover can lead to the emergence of spanning clusters. In shrublands, the effect of cover on PLR was not as high as in grasslands, and was not even significant in the model performed with our highest amount of sites (Appendix, Fig. S2). Since total cover does not differ between habitat types (F_1,68_ = 1.14, P = 0.29, in the dataset with facilitation [N=71]; F_1,109_ < 0.01, P = 0.95, in the full dataset), this result suggests either that: i) cover is not enhanced in shrublands by spatial pattern formation or ii) clumping of vegetation into large patches due to space constraints is more likely in grasslands than in shrublands. In this last case, mechanisms such as the way in which the different plant types compete (and thus repel each other) might be playing a determinant role. In shrublands, individuals are larger, and also structure forming strata that might increase competition for light that is less likely to occur between grasses. The latter is supported by the significant effect of height on PLR found in shrublands, but not in grasslands, thus indicating that the size of dominant individuals is important to define PSDs. As a result, the formation of patches in shrublands strongly relies on the size of the dominant individuals, and merging with other plants through facilitative interactions may occur in different strata, thus not showing a direct link with total cover as measured in this study.

We found that the percentage of facilitated species was an important driver of PLR in both grasslands and shrublands, whereas soil amelioration was only important in shrublands. We also found that, whereas in grasslands the number of individuals that are facilitated promoted the formation of large patches (as shown by the better fit of the PSDs to a power law), in shrublands this number had the opposite effect (Appendix, Table S1). This result suggests that, in shrublands, the number of individuals is constraining the ability of nurses to increase patch sizes, probably because the more individuals in an area, the less they can grow (Schöb et al. 2014). This result probably relates to the way in which grasses and shrubs exploit belowground resources. Grasses allow more species to coexist by expanding the niches of less-adapted species and by promoting the coexistence of understorey species through niche segregation (Soliveres et al. 2011, 2015, Berdugo et al. 2017c). We showed that this translates into the enlargement of patches. However, grasses have shallow roots. Thus, once established, beneficiaries may have problems to grow under grasses, which often compete with neighbouring plants (Paterno et al. 2016, O’Brien et al. 2017), specially at the seedling stage (Barberá et al. 2006, Soliveres et al. 2010). Therefore, the patch size would be directly related to the number of beneficiaries (which are probably small), but not to their size, in this case (Appendix, Table S1). Conversely, shrubs exploit deeper water sources, and their beneficiaries compete with each other to grow, but not with the nurse (Paterno et al. 2016, O’Brien et al. 2017), so the size of the patch is more influenced by both the size the beneficiaries and that of the nurse. The association between the percentage of facilitated species and PLR suggest that the results of previous theoretical approaches which investigated the effects of facilitation on pattern formation on models without taking into account multispecific responses of facilitation (e.g. Kéfi et al. 2007a, von Hardenberg et al. 2010) may be affected by incorporating several plant species interacting with each others. Our study highlights the necessity of including community-specific mechanisms of facilitation in models on spatial patterns. These community-specific mechanisms may depend on species pool and habitat filtering, in addition to the strength of facilitation *per se* (Fukami 2015, Berdugo et al. 2017c).

Our results relate soil amelioration mechanisms to patch formation only in shrublands, and niche-complementarity mechanisms in both grassland and shrublands. These important differences between habitat types on how facilitation drives patch formation have been previously overlooked. Both mechanisms can directly impact ecosystem functioning (through the effect of the dominants on nutrients pools, see Grime, 1998, or by enhancing the diversity of species, see Tilman et al., 1997, respectively) and are directly affected by facilitation (Maestre et al. 2003, Le Bagousse-Pinguet et al. 2014). It remains to be identified, however, if these two facilitation-related mechanisms only have an indirect effect, mediated by their effect on plant spatial patterns, or if they also have direct effects on ecosystem functioning. Future studies need to examine whether these feedbacks between more direct (soil amelioration and increases in species richness) and indirect (through the formation of spatial patterns) effects of facilitation on ecosystem functioning might feedback on each other, to better understand the overall consequences of facilitation for ecosystem functioning.

Our study also informs about the structural implications of the worldwide reported shifts from grasslands to shrublands, also known as shrub or woody encroachment (Eldridge et al. 2011). Shrub encroachment by itself does not necessarily entail losses in ecosystem diversity and/or functioning (Eldridge et al. 2011, Eldridge and Soliveres 2015). However, our findings suggest that, if the dominant growth form shifts, the processes of spatial pattern formation might be altered and become more dependent on soil amelioration. If accompanied by increases in aridity, our results predict that shrub encroachment could be associated to fewer large patches due to facilitation waning, which might be linked to functionality losses (Berdugo et al. 2017b). Indeed, the effects of shrub encroachment on soil functioning change throughout aridity gradients, shifting from positive to negative with aridity (Jackson et al. 2002, Eldridge et al. 2011, but see Knapp et al., 2008).

## CONCLUDING REMARKS

By examining the typology and drivers of PSDs in drylands worldwide, we found that PSDs tend to deviate from power laws by loosing large, but also small patches. Our results demonstrate differences in the drivers of PSDs depending on the habitat type considered. We provide evidence of the importance of positive plant-plant interactions as a driver of spatial pattern formation, an effect that was mainly due to the addition of new species to the patches, rather than by a soil amelioration effect under the nurses (although this mechanism was also important in shrublands). We also highlight the importance of plant cover and the height of the dominant species as drivers of PSDs in grasslands and shrublands, respectively. All together, this study constitutes a significant step forward on our understanding of how vegetation spatial patterns are formed and distributed in drylands, highlighting the influence of habitat-type and aridity on the relative importance of the drivers of such patterns.

## ACKNOWLEDGEMENTS

We thank Chi Xu for discussions during the processing of the images, and all the members of the EPES-BIOCOM network for the collection of field data.

## AUTHOR CONTRIBUTIONS

FTM designed the study and coordinated data collection. MB analyzed the data, helped by SK and SS. MB wrote the manuscript, with contributions from all authors.

## REFERENCES

Abades, S. R. et al. 2014. Fire, percolation thresholds and the savanna forest transition: a neutral model approach. - J. Ecol. 102: 1386–1393.

Aguiar, M. R. and Sala, O. E. 1999. Patch structure, dynamics and implications for the functioning of arid ecosystems. - Trends Ecol. Evol. 14: 273–277.

Aguiar, M. R. et al. 1992. Competition and Facilitation in the Recruitment of Seedlings in Patagonian Steppe. - Funct. Ecol. 6: 66–70.

Allington, G. R. H. and Valone, T. J. 2014. Islands of fertility: A byproduct of grazing? - Ecosystems 17: 127–141.

Anderson, J. M. and Ingram, J. S. I. 1994. Tropical soil biology and fertility: A handbook of methods. - Soil Sci. 157: 265.

Barberá, G. G. et al. 2006. Seedling recruitment in a semi-arid steppe: The role of microsite and post-dispersal seed predation. - J. Arid Environ. 67: 701–714.

Barton, K. 2016. MuMIn: Multi-Model Inference. R package version 1.15. 6.

Berdugo, M. et al. 2014. Vascular plants and biocrusts modulate how abiotic factors affect wetting and drying events in drylands. - Ecosystems 17: 1242–1256.

Berdugo, M. et al. 2017a. Dataset and R code from the article: “The interplay between facilitation and habitat type drive spatial vegetation patterns in global drylands.” - figshare. DOI:10.6084/m9.figshare.5640193

Berdugo, M. et al. 2017b. Plant spatial patterns identify alternative ecosystem multifunctionality states in global drylands. - Nat. Ecol. Evol. 1: 0003.

Berdugo, M. et al. 2017c. Species-specific adaptations determine how aridity and biotic interactions drive the assembly of dryland plant communities. - bioRxiv. DOI: https://doi.org/10.1101/147181

Bertness, M. D. and Callaway, R. 1994. Positive interactions in communities. - Trends Ecol. Evol. 9: 191–193.

Boer, M. and Puigdefábregas, J. 2005. Effects of spatially structured vegetation patterns on hillslope erosion in a semiarid Mediterranean environment: a simulation study. - Earth Surf. Process. Landforms 30: 149–167.

Bordeu, I. et al. 2016. Self-Replication of Localized Vegetation Patches in Scarce Environments. - Sci. Rep. 6: 33703.

Borthagaray, A. I. et al. 2012. Connecting landscape structure and patterns in body size distributions. - Oikos 121: 697–710.

Burnham, K. P. and Anderson, D. R. 2004. Multimodel Inference. - Sociol. Methods Res. 33: 261–304.

Cardinale, B. J. et al. 2002. Species diversity enhances ecosystem functioning through interspecific facilitation. - Nature 415: 426–429.

Clauset, A. et al. 2009. Power-law distributions in empirical data. - SIAM Rev. 51: 661–703.

Dean, W. R. J. et al. 1999. Large trees, fertile islands, and birds in arid savanna. - J. Arid Environ. 41: 61–78.

Delgado-Baquerizo, M. et al. 2013. Decoupling of soil nutrient cycles as a function of aridity in global drylands. - Nature 502: 672–676.

Eldridge, D. J. and Soliveres, S. 2015. Are shrubs really a sign of declining ecosystem function? Disentangling the myths and truths of woody encroachment in Australia. - Aust. J. Bot. 62: 594–608.

Eldridge, D. J. et al. 2011. Impacts of shrub encroachment on ecosystem structure and functioning: towards a global synthesis. - Ecol. Lett. 14: 709–722.

Fukami, T. 2015. Historical contingency in community assembly: integrating niches, species pools, and priority effects. - Annu. Rev. Ecol. Evol. Syst. 46: 1.

Goslee, S. C. et al. 2003. High-resolution images reveal rate and pattern of shrub encroachment over six decades in New Mexico, USA. - J. Arid Environ. 54: 755–767.

Grime, J. P. 1998. Benefits of plant diversity to ecosystems: immediate, filter and founder effects. - J. Ecol. 86: 902–910.

Gross, N. et al. 2017. Functional trait diversity maximizes ecosystem multifunctionality. - Nat. Ecol. Evol. 1: 0132.

Hijmans, R. J. et al. 2005. Very high resolution interpolated climate surfaces for global land areas. - Int. J. Climatol. 25: 1965–1978.

Holmgren, M. and Scheffer, M. 2010. Strong facilitation in mild environments: the stress gradient hypothesis revisited. - J. Ecol. 98: 1269–1275.

Huxman, T. E. et al. 2005. Ecohydrological Implications of Woody Plant Encroachment. - Ecology 86: 308–319.

Jackson, R. B. et al. 2002. Ecosystem carbon loss with woody plant invasion of grasslands. - Nature 418: 623–626.

Kattge, J. et al. 2011. TRY--a global database of plant traits. - Glob. Chang. Biol. 17: 2905–2935.

Kéfi, S. et al. 2007a. Spatial vegetation patterns and imminent desertification in Mediterranean arid ecosystems. - Nature 449: 213–217.

Kéfi, S. et al. 2007b. Local facilitation, bistability and transitions in arid ecosystems. - Theor. Popul. Biol. 71: 367–379.

Kettler, T. A. et al. 2001. Simplified Method for Soil Particle-Size Determination to Accompany Soil-Quality Analyses Journal Series no. 13277 of the Agric. Res. Div., Univ. of Nebraska, Lincoln, NE.. - Soil Sci. Soc. Am. J. 65: 849–852.

Knapp, A. K. et al. 2008. Shrub encroachment in North American grasslands: shifts in growth form dominance rapidly alters control of ecosystem carbon inputs. - Glob. Chang. Biol. 14:615–623.

Le Bagousse-Pinguet, Y. et al. 2014. Importance, but not intensity of plant interactions relates to species diversity under the interplay of stress and disturbance. - Oikos 123: 777–785.

Le Bagousse-Pinguet, Y. et al. 2017. Testing the environmental filtering concept in global drylands. - J. Ecol. 105: 1058–1069.

Lett, M. S. and Knapp, A. K. 2003. Consequences of shrub expansion in mesic grassland: resource alterations and graminoid responses. - J. Veg. Sci. 14: 487–496.

Liancourt, P. et al. 2017. SGH: stress or strain gradient hypothesis? Insights from an elevation gradient on the roof of the world. - Ann. Bot. 120: 29–38.

Lin, Y. et al. 2010. Spatial vegetation patterns as early signs of desertification: a case study of a desert steppe in Inner Mongolia, China. - Landsc. Ecol. 25: 1519–1527.

Ludwig, J. A. et al. 2005. Vegetation Patches and Runoff-Erosion as Interacting Ecohydrological Processes in Semiarid Landscapes. - Ecology 86: 288–297.

Lukacs, P. M. et al. 2009. Model selection bias and Freedman’s paradox. - Ann. Inst. Stat. Math. 62: 117.

Maestre, F. T. and Cortina, J. 2005. Remnant shrubs in Mediterranean semi-arid steppes: effects of shrub size, abiotic factors and species identity on understorey richness and occurrence. - Acta Oecologica 27: 161–169.

Maestre, F. T. and Escudero, A. 2009. Is the patch size distribution of vegetation a suitable indicator of desertification processes? - Ecology 90: 1729–1735.

Maestre, F. T. et al. 2001. Potential for using facilitation by grasses to establish shrubs on a semiarid degraded steppe. - Ecol. Appl. 11: 1641–1655.

Maestre, F. T. et al. 2003. Positive, Negative, and Net Effects in Grass-Shrub Interactions in Mediterranean Semiarid Grasslands. - Ecology 84: 3186–3197.

Maestre, F. T. et al. 2010. Do biotic interactions modulate ecosystem functioning along stress gradients? Insights from semi-arid plant and biological soil crust communities. - Philos. Trans. R. Soc. B Biol. Sci. 365: 2057–2070.

Maestre, F. T. et al. 2012. Plant species richness and ecosystem multifunctionality in global drylands. - Science 335: 214–218.

Manor, A. and Shnerb, N. M. 2008. Origin of Pareto-like spatial distributions in ecosystems. - Phys. Rev. Lett. 101: 268104.

Marani, M. et al. 2006. Spatial organization and ecohydrological interactions in oxygen-limited vegetation ecosystems. - Water Resour. Res. 42: 4582.

Moreno de las Heras, M. et al. 2011. Assessing landscape structure and pattern fragmentation in semiarid ecosystems using patch-size distributions. - Ecol. Appl. 21: 2793–2805.

O’Brien, M. J. et al. 2017. The shift from plant-plant facilitation to competition under severe water deficit is spatially explicit. - Ecol. Evol. 7: 2441–2448.

Ochoa-Hueso, R. et al. 2017. Soil fungal abundance and plant functional traits drive fertile island formation in global drylands. - J. Ecol. in press.

Okin, G. S. et al. 2009. Impact of feedbacks on Chihuahuan desert grasslands: Transience and metastability. - J. Geophys. Res. 114: G01004.

Paterno, G. B. et al. 2016. Species-specific facilitation, ontogenetic shifts and consequences for plant community succession. - J. Veg. Sci. 27: 606–615.

Pugnaire, F. I. et al. 1996. Facilitation and Succession under the Canopy of a Leguminous Shrub, Retama sphaerocarpa, in a Semi-Arid Environment in South-East Spain. - Oikos 76: 455–464.

Quets, J. J. et al. 2013. Unraveling landscapes with phytogenic mounds (nebkhas): An exploration of spatial pattern. - Acta Oecologica 49: 53–63.

R Development Core Team 2008. R: A Language and Environment for Statistical Computing.

Ravi, S. et al. 2008. Form and function of grass ring patterns in arid grasslands: the role of abiotic controls. - Oecologia 158: 545–555.

Rietkerk, M. et al. 2004. Self-organized patchiness and catastrophic shifts in ecosystems. - Science 305: 1926–1929.

Scanlon, T. M. et al. 2007. Positive feedbacks promote power-law clustering of Kalahari vegetation. - Nature 449: 209–212.

Schöb, C. et al. 2014. The context dependence of beneficiary feedback effects on benefactors in plant facilitation. - New Phytol. 204: 386–396.

Soliveres, S. and Maestre, F. T. 2014. Plant--plant interactions, environmental gradients and plant diversity: a global synthesis of community-level studies. - Perspect. Plant Ecol. Evol. Syst. 16: 154–163.

Soliveres, S. et al. 2010. Spatio-temporal heterogeneity in abiotic factors modulate multiple ontogenetic shifts between competition and facilitation. - Perspect. Plant Ecol. Evol. Syst. 12: 227–234.

Soliveres, S. et al. 2011. Microhabitat amelioration and reduced competition among understorey plants as drivers of facilitation across environmental gradients: Towards a unifying framework. - Perspect. Plant Ecol. Evol. Syst. 13: 247–258.

Soliveres, S. et al. 2014. Functional traits determine plant co-occurrence more than environment or evolutionary relatedness in global drylands. - Perspect. plant Ecol. Evol. Syst. 16: 164–173.

Soliveres, S. et al. 2015. A missing link between facilitation and plant species coexistence: nurses benefit generally rare species more than common ones. - J. Ecol. 103: 1183–1189.

Tilman, D. et al. 1997. The influence of functional diversity and composition on ecosystem processes. - Science 277: 1300–1302.

Tongway, D. J. et al. 2001. Banded vegetation patterning in arid and semiarid environments: ecological processes and consequences for management. - Springer Science & Business Media.

Venables, W. N. and Ripley, B. D. 2002. Modern Applied Statistics with S (Statistics and Computing). - Springer.

von Hardenberg, J. et al. 2010. Periodic versus scale-free patterns in dryland vegetation. - Proc. R. Soc. London B Biol. Sci. 277: 1771–1776.

Von Hardenberg, J. et al. 2001. Diversity of vegetation patterns and desertification. - Phys. Rev. Lett. 87: 198101.

Weltzin, J. F. and McPherson, G. R. 1999. Facilitation of conspecific seedling recruitment and shifts in temperate savanna ecotones. - Ecol. Monogr. 69: 513–534.

Wright, A. J. et al. 2017. The Overlooked Role of Facilitation in Biodiversity Experiments. - Trends Ecol. Evol. 32: 383–390.

Xu, C. et al. 2015. Can we infer plant facilitation from remote sensing? A test across global drylands. - Ecol. Appl. 25: 1456–1462.

Zomer, R. J. et al. 2008. Climate change mitigation: A spatial analysis of global land suitability for clean development mechanism afforestation and reforestation. - Agric. Ecosyst. Environ. 126: 67–80.

